# Peptide hydrophobicity and aromaticity predict multi-state translocation kinetics via protective antigen nanopores

**DOI:** 10.1101/2025.09.15.676337

**Authors:** Jennifer M. Colby, Bryan A. Krantz

**Author notes:** **Corresponding author** (BAK).

## Abstract

Single-molecule analysis of guest-host peptides translocating through the anthrax toxin protective antigen (PA) nanopore reveals a multi-state kinetic mechanism. Using K-Means clustering, four distinct conductance states, including a fully-blocked state (State 0), two intermediates (States 1 and 2), and a fully open pore (State 3) were identified. Multi-exponential kinetic analysis of state-to-state transitions was performed, and resulting lifetimes and amplitudes were correlated with molecular properties of the guest residue. Our correlation analysis of these kinetic parameters to defined molecular properties of the guest residues reveals which physical properties govern the mechanism. The fully blocked State 0 acts as a ‘hydrophobic trap,’ with the lifetime of entry transitions (e.g., 1→0) strongly predicted by side-chain hydrophobicity. Conversely, escaping this trap is a steric process governed by molecular size, though the probability of a fast escape is uniquely facilitated by aromaticity, suggesting a specific ungating interaction with the pore’s ϕ-clamp, which is consistent with clamp site dilation. Rearrangements between partially blocked states are also dominated by hydrophobicity, reflecting the side chain exploring different contacts within the pore. Final dissociation to open nanopore is a multi-pathway process where the dominant physical force depends on the starting state: escape from deeper states is an energetic battle against hydrophobicity and aromaticity, while escape from shallower states presents a final steric hurdle. Overall, this work dissects the peptide translocation process, demonstrating how distinct physical forces—hydrophobicity, sterics, and aromaticity—govern specific, sequential steps of intra-pore dynamics and release, providing a detailed energy landscape for peptide-nanopore interactions.

## Introduction

High resolution structural information (1-4) combined with dynamic translocation studies on the anthrax toxin protective antigen (PA) nanopore has allowed for elucidation of detailed molecular models of transmembrane translocation. Exploiting these structure/function approaches has made the PA nanopore a reliable biophysical model system to study protein translocation (5). As a natural protein translocase, PA is exceptionally robust and facilitates the processive, voltage- and/or proton gradient-driven transport of peptides (6, 7). It achieves this at nanomolar substrate peptide concentrations without the need for DNA tethers or other labels (8-13). PA uses multiple internal loops and clefts, called ‘peptide-clamp’ sites, such as the phenylalanine clamp (ϕ clamp), to interact non-specifically with the translocating molecule (2, 3, 9, 11, 14, 15). The highly dynamic interactions with clamp sites, notably the ϕ clamp, generate complex, multi-state ionic current signatures with distinct kinetic and conductance characteristics (9, 11-13). Deciphering these kinetic signatures is essential for mapping the underlying energy landscape of translocation, yet assigning specific physical drivers to each step of the process remains a significant challenge. Recent studies also have shown these dynamic multi-state interactions facilitate the use of the PA nanopore as a peptide biosensor platform especially when single-molecule detection is combined with powerful machine learning computational approaches (12, 13). In principle, further development of this dynamical nanopore system using high-throughput detection may enable its use as a powerful peptide sequencer— an area of research that is rapidly emerging (16-18).

Fundamental to further elucidation of either the basic science describing the molecular mechanism of translocation or the translational development of the system into a biosensor or sequencer is gaining insight into the dynamics of clamp-peptide interactions. These dynamics are rich with information for either pursuit. Nanopore engineering approaches performed to modify these dynamics systematically may produce yet more capable biosensors that are able to detect difficult analytes intractable to decipher by the unmodified wild-type nanopore (13).

Here we perform a detailed kinetic analysis of model guest-host peptide translocation via wild-type PA nanopores, examining how molecular properties of the guest residue may govern the multi-state kinetic mechanism. Specifically, we seek to disentangle the contributions of fundamental physical properties—such as side-chain hydrophobicity, sterics, and aromaticity— to the distinct kinetic phases of intra-pore rearrangement and dissociation. By systematically correlating a large set of kinetic parameters derived from multi-exponential analysis with various molecular property scales, we construct a detailed, state-dependent model of translocation. Our results demonstrate that no single property governs the entire process; rather, distinct physical forces dominate sequential, specific steps of the peptide’s journey through the pore, providing a detailed map of the underlying energy landscape.

## Results

### Multi-state kinetics of guest-host peptide translocation via PA nanopores

Previously, a large dataset of single-channel guest-host peptide translocation event streams was collected for the wild-type PA nanopore at 70 mV driving force (cis positive), pH 5.6 (12). These conditions are favorable for observing complete translocations rather than unproductive dissociations. The guest-host substrate peptide design was a 10-residue synthetic construct of sequence, KKKKKXXSXX, where X was the guest residue **(Fig. 1A)**. There were seven total peptides in the dataset. Six peptides were of pure natural stereochemistry, where X was either Ala, Leu, Phe, Thr, Trp, or Tyr; however, the seventh peptide TrpDL used a guest residue of Trp, but every other residue was ᴅ chirality instead of natural ʟ chirality. We refer to these peptides by the three-letter guest residue abbreviation, e.g., ‘guest-host Ala’ or ‘guest-host TrpDL’. The PA nanopore itself contains a central and structurally dynamic constriction site called the ϕ clamp, which forms various blocked and subconductance intermediates with translocating peptides **(Fig. 1B)**. Unlike our earlier studies employing CLAMPFIT (12), conductance state detection and labeling of the raw streams, however, was carried out using K-Means clustering with automated baseline correction (13). This more computationally efficient state labeling confirmed the presence of four conductance states in the event streams of all tested peptides: State 0 (fully blocked peptide-nanopore complex); State 1 (highly blocked but partially conducting peptide-nanopore complex); State 2 (∼50% conducting partially blocked peptide-nanopore complex); and State 3 (fully open nanopore). While these four states were observed across the guest-host peptides tested, the levels of the conductance blockade depths could vary between peptides **(Fig. 1B)**. Peptide translocation events were then detected, such that an event initiated when the nanopore transitioned from State 3 to other states, and the event terminated when the nanopore returned to State 3.

**Fig. 1.**
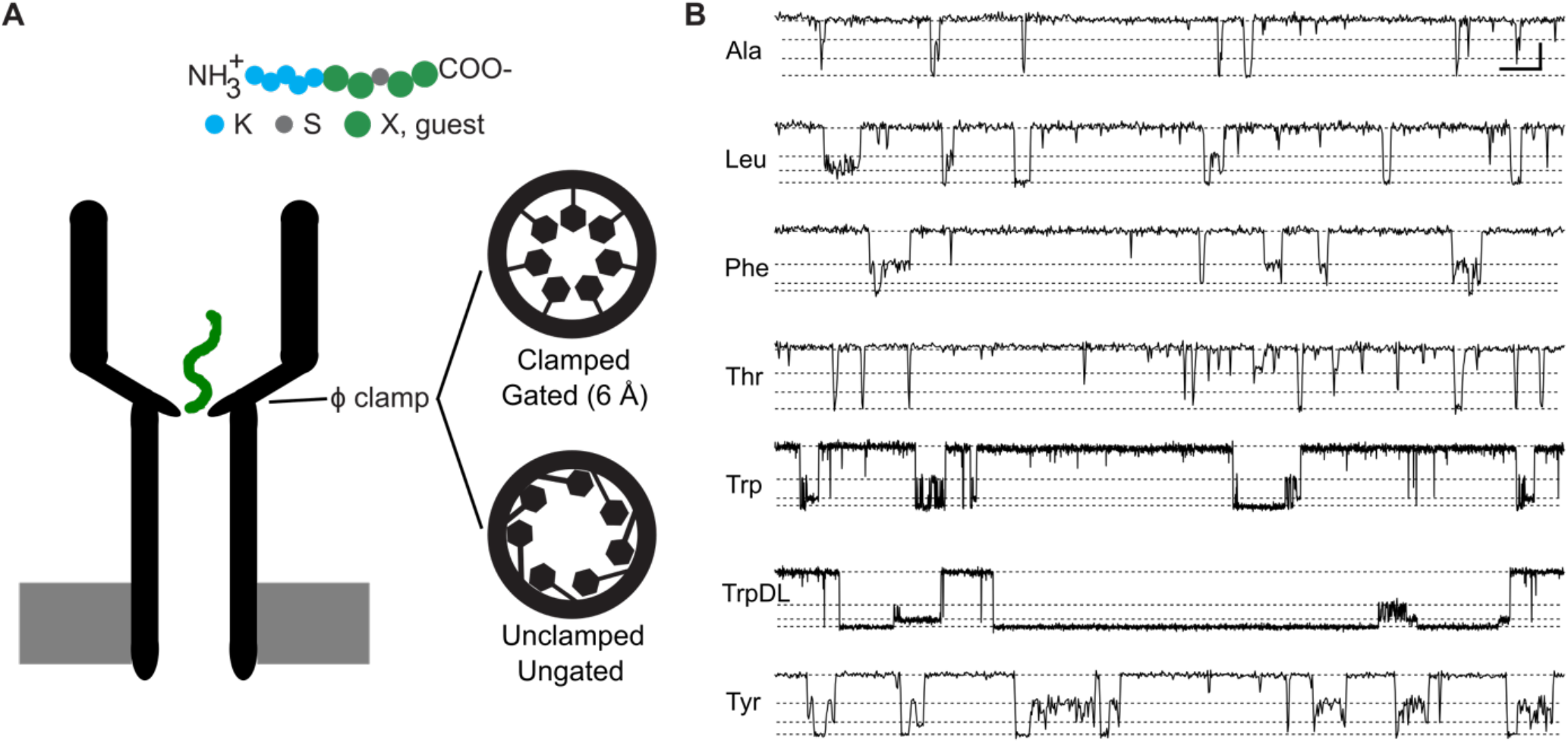
Nanopore-peptide system and translocation event streams. **(A)** Top, schematic diagram of 10-residue guest-host peptide colored by residue: Lys (blue), Ser (gray), and guest residue (green). Bottom left, schematic of the PA nanopore (black) with peptide (green) liganded at ϕ clamp constriction. Bottom right, cartoons of the structurally observed clamped and gated ϕ clamp with a 6 Å luminal diameter and of a putative unclamped, ungated state of the clamp observed electrophysiologically. **(B)** Representative recordings of guest-host peptide (20 nM) translocation event streams via wild-type PA nanopores carried out at 70 mV (cis positive) in symmetric 100 mM KCl, 20 mM succinate, 1 mM EDTA, pH 5.6. To the left are the standard three letter name for the guest residue. These nanopore-peptide systems populate multiple discrete partially or fully blocked intermediates (approximate levels indicated by dotted lines). From bottom to top of each record: fully blocked (State 0), partially blocked intermediates (State 1 and State 2), and fully open baseline (State 3). Scalebar at the upper right of the panel denotes 2 pA by 100 ms for guest-host Ala, Leu, Phe, Thr, and Tyr peptides. For guest-host Trp and TrpDL peptides, the scalebar is 2 pA by 500 ms to present longer events.

### Comprehensive kinetic analysis of multi-state kinetics

A 4×4 dwell time transition matrix was then calculated for each peptide from the entire set of translocation events. Cumulative distribution functions and corresponding survival curves were generated from the dwell times for each transition represented in the dataset. To then extract lifetimes (τ) and amplitudes (*A*), one-, two- and three-exponential decays were fitted to the natural log of the survival coordinate. Many cases deviated from linearity and exhibited two- or three-exponential relationships, indicating that there were hidden states of the system sharing the same observed conductance levels **(Fig. 2A-C)**. This result is not surprising given peptides studied previously populated a series of fully blocked intermediates (9, 10).

**Fig. 2.**
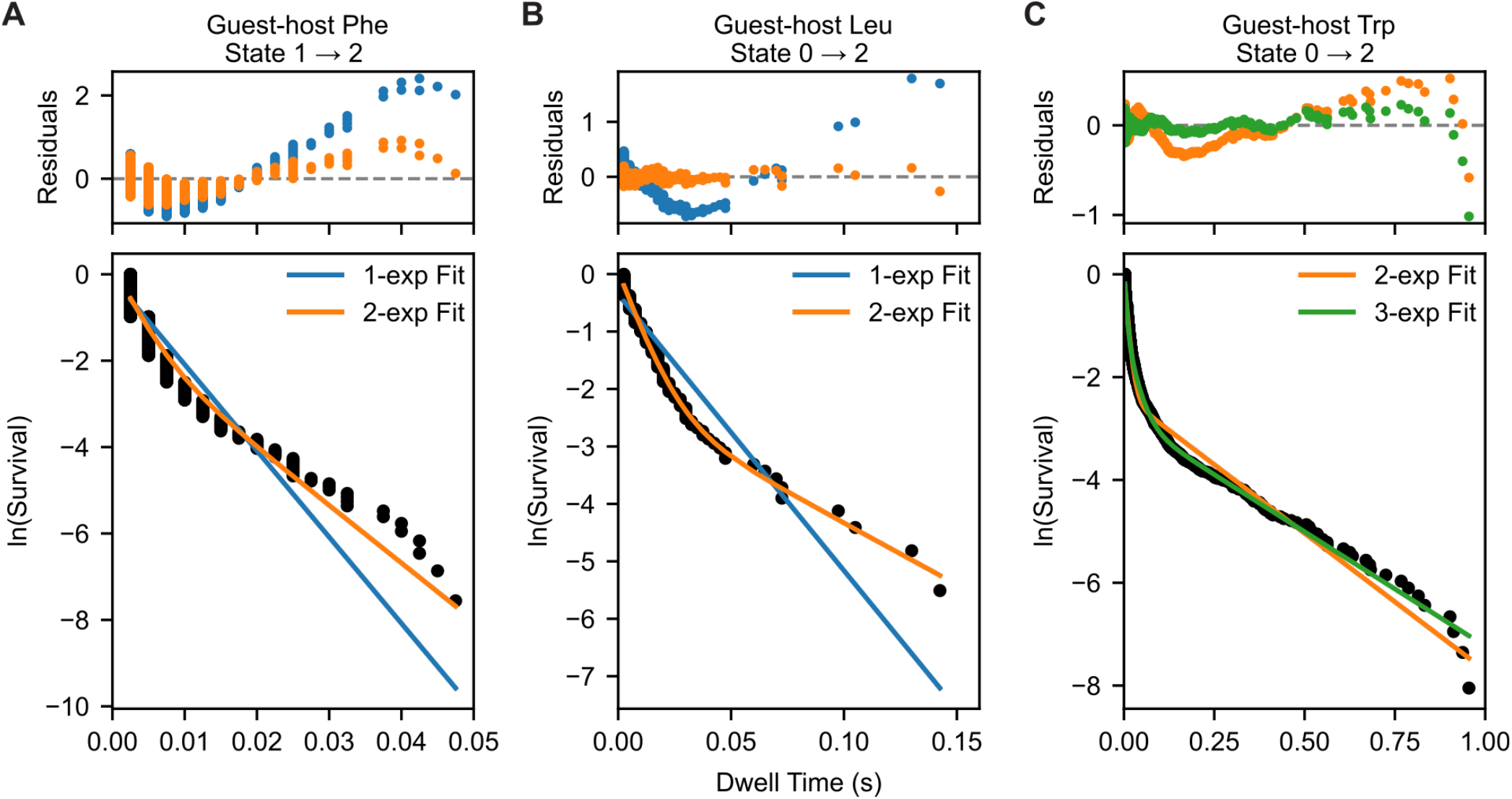
Translocation event kinetics generally fit to multi-exponential models. Natural log of the survival curves plotted against dwell time (black circles) are shown for selected transitions for the indicated peptides. Panels of selected cases are **(A)** State 1 → 2 transition for guest-host Phe peptide, **(B)** State 0 → 2 transition for guest-host Leu peptide, and **(C)** State 0 → 2 transition for guest-host Trp peptide. Exponential decay models, colored by number of exponentials (1-exp, blue; 1-exp, orange; 2-exp, green), were selected using BIC. The residuals plot at top of each panel, colored consistently with the exponential fit models, further justified these model selections.

To minimize over-fitting, the most optimal exponential decay model for each transition observed for each peptide was determined by the Bayesian Information Criterion, where a model with higher numbers of parameters was not prioritized when it failed to produce a significantly better fit. We consolidated the kinetic fit parameters for the transitions observed for each peptide as follows. A mean lifetime τ value (τ_mean_) was computed to aid in comparison of peptides when different exponential decay models were employed for a given transition. Also to aid in downstream comparative analysis the lifetimes for different exponential decay models were handled systematically by ranking as τ_fast_, τ_middle_, and τ_slow_. When the model was a one-exponential decay, then only τ_fast_ was used. When a two-exponential model was used τ_fast_ was the fasting decay, and τ_middle_ and τ_slow_ were the same second decay rate. The amplitudes were handled similarly to the τ values in this consolidation process. The entire consolidated dataset is a **Supporting Document**.

### Correlations of best kinetic fit parameters to peptide molecular properties

The consolidated kinetic parameter data for the observed transitions for each peptide were then analyzed by correlating the molecular properties of the guest residue to the base-10 log(τ) and *A* values. The properties included: molecular weight; an aromaticity Boolean; number of aromatic rings; various hydrophobicity scales (Kyte-Doolittle, Hopp-Woods, Cornette, Eisenberg, Rose, Janin, Engelman GES, Tanford, and Song); and two atomic solvation parameterized energy scales (Ooi and Krantz), where the latter, a highly specific model, included an aromatic enhancement term that was deduced previously for small molecule compound binding to the PA nanopore. The most predictive molecular properties in this set for linear correlations (based on mean *R*^2^ fit values across all transitions) were molecular weight, aromaticity, number of aromatic rings, Hopp-Woods, Tanford and Song **(Fig. 3A, Supporting Document)**. (For the limited guest amino acid residue set used, the Tanford scale was perfectly correlated with Hopp-Woods, since Hopp-Woods scale is based in part upon Tanford’s scale.) Hence solvent transfer free energies and aromatic factors were highly predictive of these guest-host peptide translocation kinetics. Probably the property of molecular weight scales well enough with these basic properties given the limited amino acid set covered in the current guest-host peptide library. It is reasonable to assume transfer free energies and aromaticity would predict kinetic parameters so well given the key interaction site, the ϕ clamp is composed of a narrow ring of seven F427 residues in the homoheptameric version of the PA nanopore, where that confined environment is hydrophobic and conducive to π-π stacking and π-dipole interactions between aromatic moieties. When we examined the best molecular property which correlates with individual kinetic parameters for the observed transitions, aromatic properties and hydrophobicity scales like Hopp-Woods or the transfer free energy scale from Song dominate **(Fig 3B; Table S1)**.

**Fig. 3.**
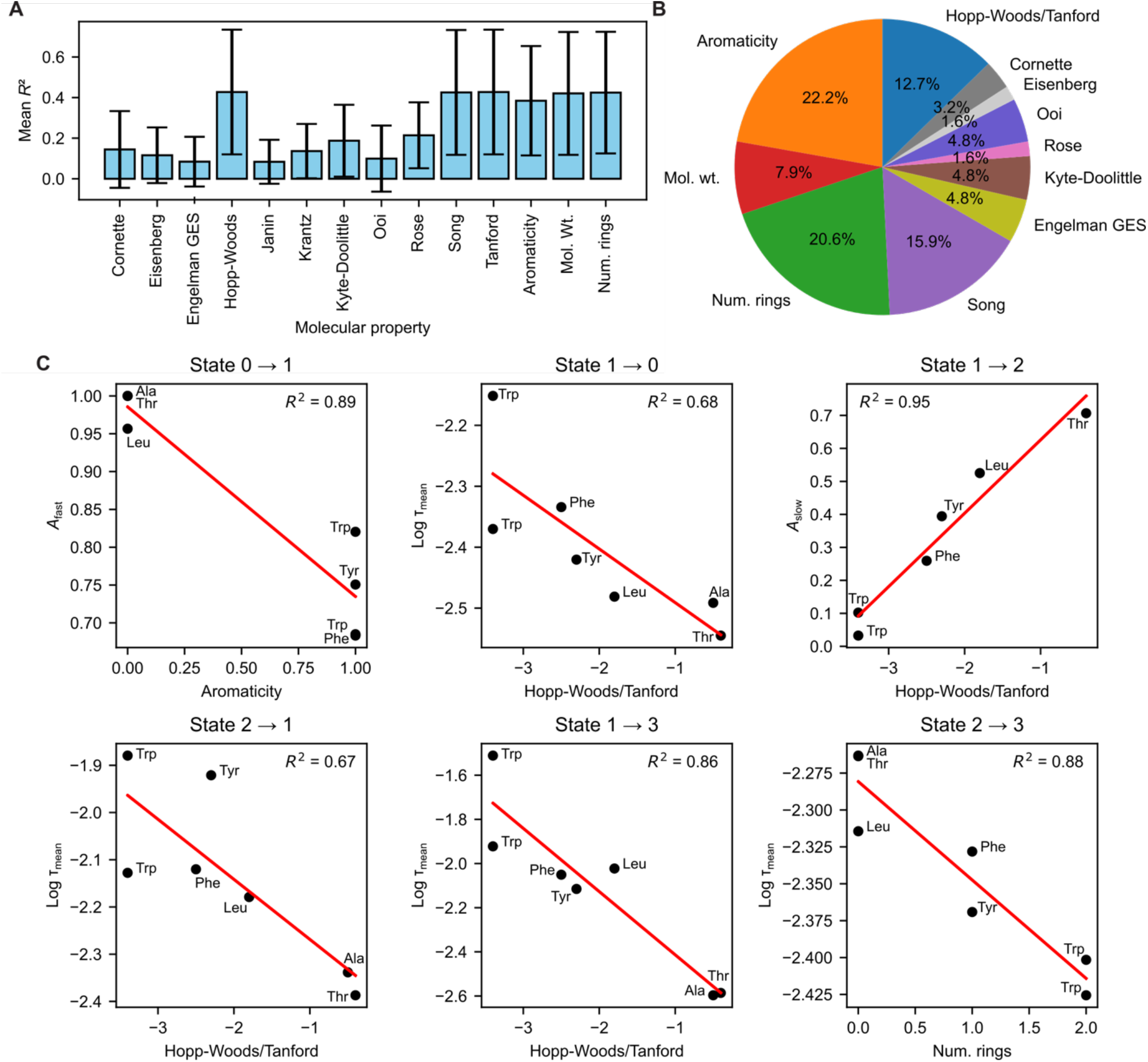
Correlation of translocation event kinetic parameters to molecular properties of guest residue. **(A)** Mean *R*^2^ for correlations of translocation event all kinetic parameters for all observed transitions versus molecular properties of the guest residue in the peptides. Error bars are the standard deviations. **(B)** Pie chart representation of the best molecular property correlation for all kinetic parameters and transitions. **(C)** Selected raw correlations of the specified kinetic parameter (y axis) to the specified molecular property (x axis) for the indicated transition (title label). The individual guest residue points are labeled by their respective 3-letter abbreviation. *R*^2^ values for the correlation are indicated. Note well there are two ‘Trp’ points for the guest-host Trp and guest-host TrpDL kinetic data.

### Molecular insights on peptide-clamp trapping and ungating

The results from our correlation of molecular properties of the guest residue to the fitted multi-state kinetic parameters highlight some well predicted steps in the overall peptide translocation mechanism. Some significant correlations to parameters among these steps are shown **(Fig. 3C)**. State 0, being fully blocked peptide bound state, represents a deep, stable binding state. The transitions into and out of this state are governed by different forces. Entering this most deeply trapped state via the State 1 → 0 transition is driven by hydrophobicity, as the lifetime for this transition, represented by log(τ_mean_), is well predicted by the Hopp-Woods/Tanford hydrophobicity scale (*R*^2^ = 0.97). Parameters for transition 2 → 0 correlate well with aromaticity and number of rings, highlighting the aromatic character of the peptide facilitates this trapping step. Therefore, State 0 is a ‘hydrophobic trap’ where the amino acid side chain has maximized its favorable, sticky contacts with the pore’s wall, likely the aromatically dense ϕ-clamp site. The more hydrophobic the residue, the longer it takes to rearrange and find this most stable state. Escaping the trap (0 → 1 and 0 → 2) is gated by aromaticity. The time it takes to move out of the deep state is best correlated with molecular weight and number of rings. The probability of making a fast exit (*A*_fast_) is best predicted by aromaticity (*R*^2^ = 0.89 for the 0 → 1 transition). Therefore, reptation out of the tight, fully-blocked trapped state is primarily a steric problem—it is a physical size challenge. However, aromatic residues seem uniquely able to initiate a rapid escape, suggesting they have access to a specific, lower-energy pathway out of the trap. In earlier work, it has been proposed that the pore may dilate at the ϕ-clamp site (9, 11) **(Fig. 1B)**, making this escape from the constricted trapped state possible; and the results herein suggest this escape is interestingly facilitated by aromaticity in the guest site of the peptide. π-π stacking and π-dipole interactions between aromatic moieties in the ϕ clamp and guest residue of the peptide may ungate the narrow configuration of the clamp to a more relaxed, dilated, or unclamped state. The most logical hypothesis is that the ∼50% conducting State 2 species is a more dilated configuration allowing significant ion leakage while engaging the peptide substrate; clearly what seems less likely is the alternative model that the ϕ clamp remains in a 6-Å luminal diameter configuration while bound to peptide and yet permits significant ion flow.

### Intermediate state dynamics in the transitions between states 1 and 2

Once outside the deepest State 0 trap, the peptide can rearrange between partially blocked states. These movements are a clear measure of the guest side chain’s ‘stickiness’. For transitions, such as 1 → 2, both the lifetimes and the amplitudes are best predicted by hydrophobicity scales, like Hopp-Woods/Tanford, with very high *R*^2^ values (e.g., *R*^2^ = 0.95 for *A*_slow_). The intermediate subconductance states (1 and 2) represent distinct conformations where the side chain makes different sets of contacts with the ϕ clamp. The energy barriers between them are directly related to the net transfer free energy of the side chain, confirming that these are hydrophobically driven rearrangements. Based on the current model, again the rearrangement of the ϕ clamp from a more constricted to a dilated configuration may underlie these interconversion steps.

### Predictive properties for the dynamics of peptide escape from the nanopore

The final step of the peptide’s translocation journey is its dissociation from the nanopore and return to the bulk solution (transitions to State 3). State 0 → State 3, while observed in the mechanism, lacks a strong correlation to the molecular properties tested. The weaker correlation suggests it’s not governed by a single, simple energy barrier. Instead, it may involve multiple, competing conformational changes in both the peptide and the pore, making the process ‘noisy’ and difficult to predict with one physical parameter. The State 1 → State 3 transition shows a much clearer and stronger signal, representing a more common route for dissociation. The mean lifetime (log τ_mean_) is best predicted by Hopp-Woods (*R*^2^ = 0.86). The slowest component (log τ_slow_) is strongly correlated with aromaticity (*R*^2^ = 0.89). Therefore, escaping from the partially blocked State 1 is an energetic battle against the ‘stickiness’ of the side chain. The rate is primarily determined by the energy required to break hydrophobic contacts. Aromatic residues face an additional, specific energy penalty to unbind, causing them to have a particularly long-lived, slow dissociation pathway from this state. Escape from the shallower trap (State 2 → State 3) shows strong correlations, but with a different set of physical properties than State 1 → State 3. The lifetime of this escape is best predicted by number of rings and the Song hydrophobicity scale. State 2 likely represents a shallower, less-engaged binding mode of a potentially dilated clamp. The strong correlation with number of rings suggests the final step is a steric or size-dependent hurdle. The bulk and shape of the side chain, particularly the extensive surface area of aromatic rings, dictate how easily it can navigate the final exit from the nanopore. A summary of the molecular interactions defining these many kinetic transitions is presented in **Fig. 4**.

**Fig. 4.**
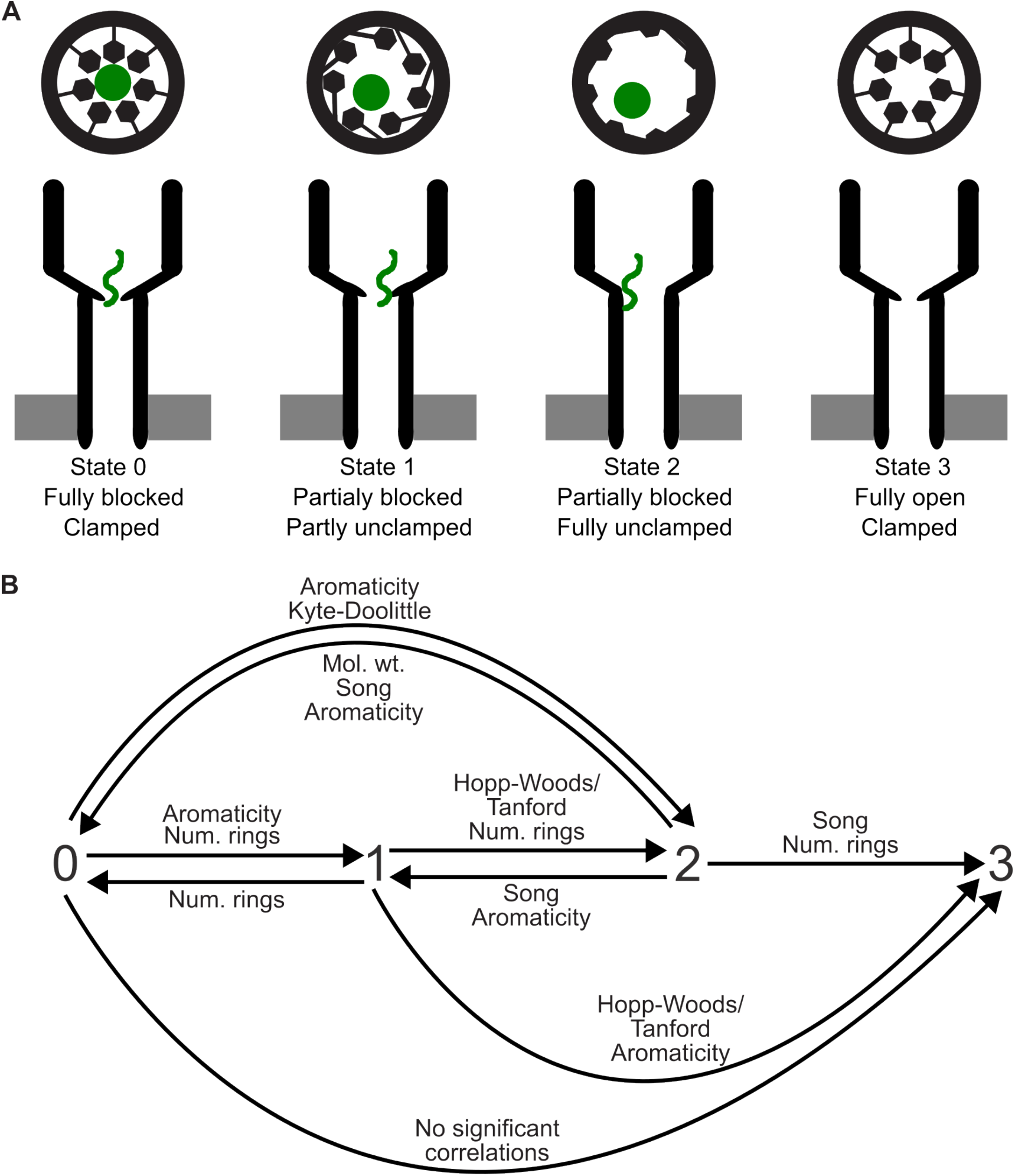
Models of polypeptide translocation via PA nanopores. **(A)** Cartoons of the discrete enumerated conductance state intermediates may reveal differential discrete gated states of the ϕ-clamp site (upper diagram) where the peptide ligand is the green circle. PA nanopores (lower diagram) show small peptides (green) can be trapped in intermediates that partially conduct due to clamp dilation. **(B)** Kinetic transition flow diagram describing translocation events, where labeled guest residue molecular properties most significantly correlate with kinetic parameters for the indicated transition. This diagram is simplified since kinetic analysis **(Fig. 2)** reveals many transitions are multi-exponential, indicating other hidden states with identical conductance levels are populated during translocation.

## Discussion

### Deconstruction of multi-state translocation kinetics by determining governing physical forces

This study successfully deconstructed the complex, multi-state kinetics of guest-host peptide translocation through the PA nanopore. The kinetic parameters (lifetimes and amplitudes) for distinct state transitions are not in fact random but are strongly and predictably correlated with fundamental molecular properties of the guest residue in the translocated peptide. This is not surprising given that peptides have high affinity for the ϕ-clamp site, which is a narrow ring of seven F427 residues in the nanopore (1, 3, 4, 14). Given this hydrophobic and aromatically dense molecular environment we surveyed correlations to a variety of relevant molecular properties of the guest residue in the peptide. Side chain transfer free energy scales (like Hopp-Woods/Tanford and Song), molecular weight, and aromaticity/number of rings were the most powerful predictors, far outperforming other hydrophobicity scales and even more complex atomic solvation parameterized models for this system. Furthermore, our proposed physical model of these events also includes dynamic state changes in the ϕ clamp structure, where leaky subconductance intermediates are proposed to be more dilated conformers of the clamp site. These peptide-clamp dynamics and their molecular preferences will not only provide further insight on the biophysical mechanism of translocation (5) but also facilitate more translational nanopore biosensor development (12, 13).

### ϕ clamp is a dynamic and gated translocase active site

The multi-state peptide translocation kinetic model, observation of subconductance states, and high degree of correlation of kinetic parameters to aromaticity and transfer free energies implies the structure of the ϕ clamp may be dynamic **(Fig. 4A)**. The strong correlations with these side-chain molecular properties throughout the translocation mechanism **(Fig. 4B)** for the three peptide-bound conductance states strongly implicate the clamp as the primary interaction site in the nanopore. These data support the hypothesis that a single static, 6-Å luminal diameter ϕ clamp conformation cannot explain the multiple, distinct, and partially conducting states (especially State 2), which is somehow peptide occupied but ∼50% conducting. Instead, these kinetic peptide translocation data are best explained by a model where the ϕ clamp itself is dynamic, adopting differentially ‘gated’ states (clamped/constricted vs. unclamped/dilated) in response to its interactions with the peptide.

Other prior studies lend support to this model. Tetraalkylammonium sizing ions can translocate via the nanopore, estimating the narrowest dimension at 11 Å (19). Two binding models of the ϕ clamp (called ‘clamped’ and ‘unclamped’) were demonstrated to be under allosteric control (9). A large Trp-rich stapled-helix was shown to be able to translocate, albeit more slowly than an unstapled control (11). Structurally, however, only the narrowest 6-Å luminal diameter configuration has been observed by cryo-electron microscopy (1, 3, 4). Moreover, other studies have argued against a dilated ϕ clamp configuration, albeit those analyses neither used polypeptide substrates nor probed the partially conducting intermediates (20).

The findings made here that aromaticity relates to escape from the deepest trap (0→1 and 0→2 transitions) provide new key mechanistic evidence on this dynamic clamp model. We now propose specific π-π and/or π-dipole interactions between the guest residues and the F427 residues can actively modulate conformational changes (dilation) in the clamp, which controls the energy barrier for progression of the peptide translocation process.

### State-dependent mechanistic model of peptide-clamp dynamics

The results here support a detailed mechanistic model, where different physical forces dominate distinct phases of the peptide’s journey through the PA nanopore **(Fig. 4)**. While there is no one single pathway through the pore as evidenced by the web-like kinetic flow diagram **(Fig. 4B)**, we have simplified discussion here by focusing on some key stages.

A central feature of this mechanism is the role of State 0 as a deep, kinetically stable ‘hydrophobic trap,’ which likely represents the guest residue’s maximal interaction with the non-polar ϕ-clamp site. The evidence for this is exceptionally strong: the lifetime to enter this state via the 1→0 transition is almost perfectly predicted by the guest residue’s hydrophobicity, as measured by the Hopp-Woods/Tanford transfer free energy scale (R^2^ = 0.97). This indicates a hydrophobically-driven process where the side chain settles into its most energetically favorable conformation within the clamp. In contrast, escaping this deep trap is governed by a fundamentally different set of forces. The lifetimes of the 0→1 and 0→2 transitions are best correlated with steric factors like molecular weight and number of aromatic rings, suggesting reptation out of the constricted state is a physical size challenge. Critically, the probability of a fast escape is gated by aromaticity (R^2^ = 0.89 for *A*_fast_ in the 0→1 transition), implying that aromatic residues have a unique ability to initiate a rapid exit.

Once outside the deepest trap, the peptide explores intermediate, partially blocked conformations (States 1 and 2). The transitions between these states represent a dynamic ‘hydrophobic dance’ within the pore. The kinetics of these rearrangements (e.g., 1 ↔ 2) are best correlated with transfer free energy scales, confirming that they are driven by the making and breaking of different sets of non-covalent contacts between the peptide side chain and the ϕ-clamp. These intermediate states are not merely transient steps but represent distinct, energetically accessible substates on the translocation pathway, the relative populations of which are dictated by the guest residue’s ‘stickiness.’

### Limitations and future directions

While the earliest work on the ϕ clamp considered the site functioning more statically and a simpler binding interaction was described (14), these new data lead to a high-resolution kinetic model, revealing the dynamic, multi-step nature of the peptide-clamp interactions. The kinetic scheme elaborated here **(Fig. 4)** is, nonetheless, oversimplified since there are still many remaining hidden states with overlapping conductance levels given the multi-exponential kinetics observed **(Fig. 2)**. Moreover, undoubtedly the guest-host residue construct and amino acid series tested herein is limited, and future work needs to examine a more broadened selection of residues and utilize natural sequences with more nuanced characteristics and compositions. This study also focused on a single voltage and pH condition, albeit the goal was to explore a strong and constant translocation promoting condition. Largely natural peptide backbone constructs were tested, although one guest-host peptide, TrpDL, explored mixed backbone stereochemistry.

With these limitations in mind, this work provides a clear roadmap for future work. We can use site-directed mutagenesis to engineer the pore (e.g., F427A) and reassess the kinetic dependencies on molecular properties, such as aromaticity and transfer free energies, since obviously the key ϕ-clamp moiety in that loop will have been ablated. Structural studies are needed to investigate alternate ϕ clamp conformations, especially in peptide liganded states, which promote subconductance state intermediates **(Fig. 1B)**. Alternatively, molecular dynamics simulations may be used to better visualize the proposed peptide conformations and clamp movements. Finally, it will be important to expand the chemical space of the guest-host peptide series. Peptides with charged (Asp, Arg, His, Lys), hydrophilic (Asn, Gln, Ser), or more uniquely shaped (Pro) guest residues can extend the limits of the current model. Future studies on natural peptide sequences are needed to determine how general the observed intermediates are and to discover more nuanced rules for clamp-peptide dynamics. These types of experiments may play well into exploring the nanopore as a viable sequencer.

## Conclusion

### Significance of findings

This study provides a comprehensive kinetic map of peptide translocation, revealing a state-dependent interplay of fundamental physical forces. While four conductance states are reported for this nanopore system, many hidden states were successfully detected, echoing earlier work on longer peptide translocation, which is characterized by a complex series of states of fully blocked conductance levels. Aromaticity/number of rings and transfer free energy dependencies dominate this kinetic landscape, lending further support that the observed multi-step process involves many interactions with and re-arrangements of the aromatic and hydrophobic ϕ clamp.

### Broader impact on peptide biosensor development

The detailed kinetic map of peptide-clamp interactions presented here provides more than just a fundamental understanding of the PA translocation mechanism; it offers a blueprint for advancing the translational applications of this nanopore system. The multi-state current blockades are not random noise but are, in fact, information-rich signals highly sensitive to the molecular properties of individual amino acid side chains. Each phase of translocation—from the initial capture and deep trapping to intra-pore rearrangements and final dissociation—is governed by a distinct set of physical forces, creating a complex kinetic ‘fingerprint’ for each guest-host peptide. This inherent sensitivity is precisely what can be exploited in next-generation nanopore biosensors. Machine learning algorithms, including the use of deep neural networks, are exceptionally well-suited to learn and classify the subtle patterns within these dynamic, multi-state translocation events (12, 13, 21). The finding that specific steps are predictably governed by properties like hydrophobicity, size, and aromaticity suggests that a machine learning model could be trained to infer these properties from the electrical signal alone. By understanding the underlying physics, we can better engineer the feature extraction process for these models, moving beyond a ‘black box’ approach. Ultimately, this detailed mechanistic insight is a critical step toward the rational design of the PA nanopore system for high-fidelity peptide sensing and, potentially, the long-term goal of single-molecule peptide sequencing.

## Materials and Methods

### Nanopore guest-host peptide system and data acquisition

The wet lab procedures and the collection of raw datasets used in this analysis were previously described (12). Briefly, monomeric 83-kDa PA (PA_83_) preprotein and their homoheptameric prepore oligomers (PA_7_) were produced as described (14, 22). Ten-residue guest-host peptides of the general sequence, KKKKKXXSXX, where X = A, L, F, T, W, and Y, were synthesized with standard ʟ amino acids (8, 11-13, 21) (Elim Biopharmaceuticals). One stereochemical variant of X = W (called TrpDL) was produced, where instead of synthesizing the peptide with uniform ʟ amino acids, an alternating pattern of ᴅ and ʟ amino acids was used.

Planar lipid bilayers were used to acquire peptide translocation event streams via PA nanopores using an Axopatch 200B amplifier system (Molecular Devices) as previously described (11, 12, 22, 23). The basic experimental conditions used in peptide translocation recordings were 100 mM KCl, 20 mM succinic acid, 1 mM EDTA, pH 5.6 under a constant +70 mV driving force (cis positive). A 400 Hz sampled dataset was used, where in some cases higher time sampled raw data were downsampled to 400 Hz. The previously deposited CLAMPFIT-labeled data were, however, re-labeled for conductance state using an updated more computationally efficient K-Means clustering routine, as described below (13).

### Hardware and software used for stream preprocessing and analysis

Anaconda was used to create a Python 3.10.16 environment, where other standard modules were installed.

The hardware used in processing and analysis was a 2025 MacBook Pro with M4 Apple Silicon and 24 GB of RAM. All source code is available at GitHub (https://github.com/bakrantz/Pept-Class).

### Conductance state labeling of raw peptide translocation event streams

Four discrete conductance states were detected in translocation event stream recordings using K-Means clustering, where baseline current drift was corrected by applying a moving window average offset (4000 time point window) (13). While rare additional states were noted, the data were labeled for the four dominant states. By convention, the fully blocked peptide-bound state was State 0, the intermediate closest to the fully blocked state was State 1, the intermediate closest to the open state was State 2, and the open state was State 3. Ultimately, the state labeling produced a three-column CSV file of the stream with columns ‘Time’, ‘Current’, and ‘State’. >90% of time points were labeled consistently when comparing our K-Means algorithm and the more traditional CLAMPFIT. All labeled CSV stream files for the seven peptides were entered into a local annotated peptide database to aid loading large numbers of individual streams.

### Event segmentation and kinetic analysis

Raw state-labeled event streams were segmented into translocation events, as described (12). The minimum event duration, which served as an effective filter for excluding very short events which can be unrelated to peptide translocation events, was set to 5 ms to ignore the shortest 2.5 ms wide spike events (single time point at 400 Hz sampling). Each segmented event was defined as initiating when the current changed from the fully open state (State 3, corresponding to baseline current) to any peptide-bound state (State 0, 1, or 2) and terminating when the current returned to State 3. From these segmented events, the corresponding state sequences were extracted.

A comprehensive list of dwell times, *t*, for each observed transition was computed in a matrix arrangement (‘from state’ as columns by ‘to state’ as rows) from the segmented translocation event state sequences observed for a given peptide. A cumulative distribution function (CDF), survival curve, *S* (1 - CDF), and natural log of the survival curve, ln(*S*), were determined for each position in the transition matrix. Different exponential decay models were fitted to ln(*S*), including single- (Eq. 1), double- (Eq. 2), and triple-exponential decay functions (Eq. 3), yielding respective lifetimes, τ, and amplitudes, *A*.

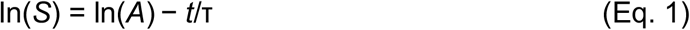

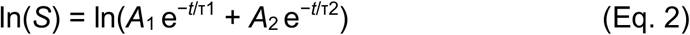

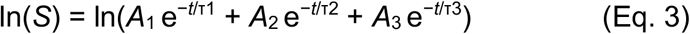

To enable comparison of complex multi-exponential kinetic transitions to single-exponential ones, mean lifetime (T_mean_) was calculated by T_mean_ = ∑ T_i_*A*_i_.

### Kinetic model selection and parameter consolidation

The best model for each transition in the case of each peptide was determined by the Bayesian Information Criterion (BIC), which penalizes over parameterization. In terms of the residual sum of squares (RSS) from the respective curve fits, BIC = *n* ln(RSS/*n*) + *k* ln(*n*). The best exponential decay model was then selected based on the lowest BIC, ignoring, of course, all fits that failed to converge. These selections were confirmed to be robust by applying a more stringent ΔBIC threshold of 6, which resulted in an identical model choice for all but one of the transitions studied, with no significant impact on the final correlation analysis. To aid in downstream comparative analysis the lifetimes for different exponential decay models were handled systematically by ranking as τ_fast_, τ_middle_, and τ_slow_. When the model was a one-exponential decay, then only τ_fast_ was used. When a two-exponential model was used τ_fast_ was the fastest decay, and τ_middle_ and τ_slow_ were the same second, slower decay lifetime. The respective fast, middle and slow amplitudes were handled similarly as the τ values in this consolidation process.

### Correlation of kinetic parameters and guest residue molecular properties

The consolidated kinetic parameter data for the observed transitions for each peptide were then analyzed by correlating the molecular properties of the guest residue to the base-10 log(τ) and raw *A* values, including τ_mean_, τ_fast_, τ_middle_, τ_slow_, *A*_fast_, *A*_middle_, and *A*_slow_. The molecular properties of the guest residue included in this analysis were molecular weight; an aromaticity Boolean; number of aromatic rings; various hydrophobicity scales (Kyte-Doolittle (24), Hopp-Woods (25), Cornette (26), Eisenberg (27), Rose (28), Janin (29), Engelman GES (30), Tanford (31), and Song (32)); and two atomic solvation parameterized (ASP) energy scales (Ooi (33) and Krantz (14)), where the latter, a highly specific model, included an aromatic enhancement term (0.7 kcal/mol/aromatic ring) that was deduced previously for small molecule compound binding to the PA nanopore. For the Ooi and Krantz ASP-based scales, solvent accessible surface areas (SASA) were computed on simple models of guest residues generated in CHIMERA (34) using GETAREA (35); their solvation energies were computed from SASA data organized by atom type using values taken from Ooi (33). Linear regression analysis was performed between the kinetic parameters and molecular properties for each transition observed. The best molecular property for each observed kinetic transition was determined in the analysis **(Table S1)**. The best overall molecular properties, which were most predictive of all kinetic parameters over all transitions, were determined by mean *R*^2^ values over all parameters for all observed kinetic transitions.

## Data availability statement

All experimental electrophysiological records (with K-Means state labeling) and related source code are publicly available. The datasets used in this manuscript have been deposited in the Zenodo repository under the DOI: xxxx. The source code is maintained on a GitHub repository (https://github.com/bakrantz/Pept-Class).

## Acknowledgments

We thank members of the department for useful feedback and discussions. J.M.C. and B.A.K. conceived of the experiments. J.M.C. collected the data. J.M.C. and B.A.K performed analyses. B.A.K. and J.M.C. wrote the manuscript. Portions of this document, including some of the Perl and Python code and language refinement, were generated with the assistance of AI-powered tools. All content was reviewed and approved by the authors, who take full responsibility for its accuracy.

## Notes

### Competing Interest Statement

The authors have declared no competing interest.

### Funding

- National Institutes of Health: 1R01AI077703, 5R21AI124020

