## Supplemental Table 1 for "Peptide hydrophobicity and aromaticity predict multi-state translocation kinetics via protective antigen nanopores"

**Table S1. Best correlating molecular properties per kinetic parameter per transition.**

| Transition | Kinetic parameter | Best molecular property | Best $R^2$ | Best *p* value | *n* |
|---|---|---|---|---|---|
| 0→1 | A_fast | Aromaticity | 0.8852 | 0.0016 | 7 |
| 0→1 | log_tau_middle | Mol. wt. | 0.7623 | 0.0103 | 7 |
| 0→1 | log_tau_mean | Num. rings | 0.6781 | 0.0228 | 7 |
| 0→1 | log_tau_fast | Song | 0.5191 | 0.0678 | 7 |
| 0→1 | A_middle | Engelman GES | 0.3649 | 0.1509 | 7 |
| 0→1 | log_tau_slow | Engelman GES | 0.211 | 0.2998 | 7 |
| 0→1 | A_slow | Aromaticity | 0.1619 | 0.3708 | 7 |
| 0→2 | A_fast | Aromaticity | 0.8019 | 0.0064 | 7 |
| 0→2 | log_tau_slow | Kyte-Doolittle | 0.7402 | 0.013 | 7 |
| 0→2 | A_slow | Rose | 0.5667 | 0.0508 | 7 |
| 0→2 | log_tau_mean | Num. rings | 0.5546 | 0.0548 | 7 |
| 0→2 | log_tau_middle | Song | 0.5392 | 0.0602 | 7 |
| 0→2 | A_middle | Aromaticity | 0.312 | 0.1925 | 7 |
| 0→2 | log_tau_fast | Aromaticity | 0.1693 | 0.3591 | 7 |
| 0→3 | A_middle | Ooi | 0.7359 | 0.0135 | 7 |
| 0→3 | log_tau_slow | Kyte-Doolittle | 0.6579 | 0.0268 | 7 |
| 0→3 | A_fast | Ooi | 0.5643 | 0.0516 | 7 |
| 0→3 | log_tau_mean | Num. rings | 0.4681 | 0.09 | 7 |
| 0→3 | A_slow | Num. rings | 0.3515 | 0.1606 | 7 |
| 0→3 | log_tau_middle | Num. rings | 0.3175 | 0.1878 | 7 |
| 0→3 | log_tau_fast | Engelman GES | 0.2288 | 0.2776 | 7 |
| 1→0 | log_tau_slow | Num. rings | 0.8502 | 0.0031 | 7 |
| 1→0 | log_tau_mean | Num. rings | 0.7097 | 0.0174 | 7 |
| 1→0 | A_fast | Eisenberg | 0.5277 | 0.0645 | 7 |
| 1→0 | A_slow | Kyte-Doolittle | 0.4752 | 0.0867 | 7 |
| 1→0 | log_tau_middle | Num. rings | 0.2806 | 0.2214 | 7 |
| 1→0 | A_middle | Ooi | 0.2128 | 0.2974 | 7 |
| 1→0 | log_tau_fast | Aromaticity | 0.1829 | 0.3386 | 7 |

| Transition | Kinetic parameter | Best molecular property | Best $R^2$ | Best *p* value | *n* |
|---|---|---|---|---|---|
| 1→2 | A_slow | Hopp-Woods | 0.949 | 0.001 | 6 |
| 1→2 | log_tau_mean | Hopp-Woods | 0.8583 | 0.0027 | 7 |
| 1→2 | log_tau_slow | Num. rings | 0.8538 | 0.0084 | 6 |
| 1→2 | log_tau_middle | Tanford | 0.5316 | 0.1001 | 6 |
| 1→2 | log_tau_fast | Hopp-Woods | 0.513 | 0.0702 | 7 |
| 1→2 | A_fast | Mol. wt. | 0.4375 | 0.1056 | 7 |
| 1→2 | A_middle | Aromaticity | 0.3014 | 0.2592 | 6 |
| 1→3 | log_tau_middle | Hopp-Woods | 0.927 | 0.0086 | 5 |
| 1→3 | log_tau_slow | Aromaticity | 0.8919 | 0.0156 | 5 |
| 1→3 | log_tau_mean | Hopp-Woods | 0.8591 | 0.0027 | 7 |
| 1→3 | log_tau_fast | Cornette | 0.8263 | 0.0046 | 7 |
| 1→3 | A_middle | Hopp-Woods | 0.6775 | 0.0869 | 5 |
| 1→3 | A_slow | Tanford | 0.5327 | 0.1616 | 5 |
| 1→3 | A_fast | Song | 0.3926 | 0.1322 | 7 |
| 2→0 | A_slow | Mol. wt. | 0.9968 | 0.0001 | 5 |
| 2→0 | log_tau_slow | Song | 0.9554 | 0.0041 | 5 |
| 2→0 | log_tau_middle | Aromaticity | 0.8071 | 0.0383 | 5 |
| 2→0 | log_tau_mean | Aromaticity | 0.7263 | 0.0149 | 7 |
| 2→0 | A_fast | Num. rings | 0.5446 | 0.0583 | 7 |
| 2→0 | A_middle | Aromaticity | 0.4034 | 0.2496 | 5 |
| 2→0 | log_tau_fast | Cornette | 0.1644 | 0.3669 | 7 |
| 2→1 | log_tau_fast | Aromaticity | 0.9238 | 0.0006 | 7 |
| 2→1 | A_slow | Aromaticity | 0.7722 | 0.0092 | 7 |
| 2→1 | log_tau_mean | Song | 0.7659 | 0.0099 | 7 |
| 2→1 | log_tau_middle | Aromaticity | 0.6899 | 0.0207 | 7 |
| 2→1 | log_tau_slow | Mol. wt. | 0.5786 | 0.0471 | 7 |
| 2→1 | A_fast | Song | 0.5752 | 0.0481 | 7 |
| 2→1 | A_middle | Song | 0.1674 | 0.3621 | 7 |
| 2→3 | log_tau_mean | Song | 0.9401 | 0.0003 | 7 |

| Transition | Kinetic parameter | Best molecular property | Best $R^2$ | Best *p* value | *n* |
|---|---|---|---|---|---|
| 2→3 | A_middle | Song | 0.912 | 0.0008 | 7 |
| 2→3 | A_slow | Song | 0.912 | 0.0008 | 7 |
| 2→3 | log_tau_middle | Num. rings | 0.8369 | 0.0039 | 7 |
| 2→3 | log_tau_slow | Num. rings | 0.8369 | 0.0039 | 7 |
| 2→3 | log_tau_fast | Num. rings | 0.6807 | 0.0223 | 7 |
| 2→3 | A_fast | Mol. wt. | 0 | 1 | 7 |
